# Insights into global antimicrobial resistance dynamics through the sequencing of enteric bacteria from U.S. international travelers

**DOI:** 10.1101/2025.01.27.635056

**Authors:** Sushmita Sridhar, Colin J Worby, Ryan A Bronson, Sarah E Turbett, Elizabeth H Oliver, Terrance Shea, Sowmya R Rao, Vanessa Sanchez, Margaret V Becker, Lucyna Kogut, Damien Slater, Jason B Harris, Maroya Spalding Walters, Allison Taylor Walker, Mark C Knouse, Daniel T Leung, Paul Kelly, Edward T Ryan, Regina C LaRocque, Ashlee M Earl

## Abstract

Antimicrobial resistance (AMR) is an urgent threat to public health, but gaps in surveillance limit the detection of emergent novel threats and knowledge about the global distribution of AMR genes. International travelers frequently acquire AMR organisms, and thus may provide a window into AMR dynamics in otherwise poorly monitored regions and environments. To assess the utility of travelers as global AMR sentinels, we collected pre- and post-travel stool samples from 608 travelers, which were screened for the presence of extended-spectrum beta-lactamase producing Enterobacterales, carbapenem-resistant Enterobacterales, and *mcr*-mediated colistin-resistant Enterobacterales. A total of 307 distinct AMR organisms were sequenced in order to determine genotypic patterns and their association with travel region and behavior. Travel-associated AMR organisms were overwhelmingly *E. coli*, which exhibited considerable phylogenetic diversity regardless of travel region. However, the prevalence of resistance genes varied by region, with *bla*_CTX-M-55_ and *bla*_CTX-M-27_ significantly more common in travelers returning from South America and South-Eastern Asia, respectively. Hybrid assembly and plasmid reconstruction revealed the genomic neighborhood of *bla*_CTX-M-55_ frequently matched a motif previously linked to animal populations. Contact with animals was also associated with virulence factors in acquired AMR organisms, including carriage of the ColV plasmid, a driver of avian pathogenic *E. coli*. We identified novel variants of the *mcr-1* gene in strains acquired from Western Africa, highlighting the potential for traveler surveillance to detect emerging clinical threats. Ongoing efforts to track travel-acquired organisms could complement existing global AMR surveillance frameworks.

**Importance:** We collected pre- and post-travel stool samples from 608 U.S. international travelers, which were screened for the presence of three clinically relevant antimicrobial-resistant organisms, with the goal of understanding the utility of travelers as sentinels for emerging AMR and identifying regional resistance patterns. We sequenced 307 organisms and uncovered significant diversity and geographic heterogeneity in AMR organisms. We found significant differences in certain resistance gene alleles of travelers returning from South America and South-Eastern Asia, traveler contact with animals associated with virulence factors, including a plasmid that is a known driver of avian pathogenic *E. coli*, and novel variants of the colistin resistance *mcr-1* gene in strains acquired from Western Africa. This work emphasizes the value of monitoring international travelers as a proxy for understanding AMR patterns globally and could be an important addition to existing global AMR surveillance frameworks.

## Introduction

Antimicrobial resistance (AMR) is a pressing global health concern, with recent estimates showing that approximately 5 million deaths in 2019 were associated with drug resistant infections (1). One of the drivers of AMR is overuse of antimicrobials in the hospital and community, which spill over into the environment (2, 3). Global differences in antibiotic administration and bacterial transmission rates have contributed to a highly heterogeneous AMR burden worldwide, with higher rates of AMR infections in many lower- and middle-income countries (LMIC), and rates are predicted to increase (4); (1). Extended-spectrum beta-lactamase-producing Enterobacterales (ESBL-PE) are considered particularly problematic, as over 60% of antimicrobials used are beta-lactams (5, 6). The ability to detect the emergence and spread of novel AMR threats is hampered by limited surveillance in many parts of the world (7) with genomic surveillance in particular predominantly occurring in healthcare settings in wealthy countries.

International travel is a risk factor for acquisition of AMR organisms (AMROs) (8) and plays a role in the global spread of antibiotic resistance genes (9). However, several recent studies of international travelers have identified ESBL-PE acquisition rates of up to 60%, depending on travel destination and traveler behavior (10–12); (13). By inadvertently sampling from microbial reservoirs, including food, water and environmental sources in diverse global destinations, travelers may act as sentinels for the emergence and spread of novel AMR threats. Genomic surveillance of travelers could thus provide a proxy insight into global AMR dynamics.

Investigations to date of ESBL-PE from travelers have typically involved culturing organisms from stool on selective media to profile for specific AMR, followed by targeted PCR-based assessment of genetic resistance factors such as beta-lactamase and carbapenemase genes (12, 14–16). Few studies have explored in detail the genetic features and phylogenetic patterns of travel-acquired multidrug resistant (MDR) Enterobacterales that might be relevant for understanding dissemination of novel AMR genes and virulence factors (17, 18). While some recent studies have sequenced isolates from travelers, these have been small in scale (19–21), focused on specific travel destinations (18, 22, 23), or been restricted to symptomatic individuals post-travel (24–27).

Previously, in an untargeted metagenomic analysis of stool samples from U.S. international travelers, we identified widespread acquisition of *E. coli* strains across a range of destinations, as well as a significant increase in the prevalence of AMR genes (28). However, due to the complexity and limited resolution of metagenomic sequence data, we were unable to identify specific AMR gene alleles with nucleotide-level accuracy, or to link genes with their plasmid or chromosomal hosts; such data are essential to tracking AMR gene spread via clonal dissemination or horizontal gene transfer. In this study, we sequenced and analyzed 307 AMRO isolates across three resistance phenotypes (ESBL-PE, carbapenem-resistant Enterobacterales (CRE), and *mcr*-mediated colistin-resistant Enterobacterales (mcr-E)) collected before and after international travel from our previously described cohort of U.S. travelers (28). In this study we sought to assess the utility of travelers as ‘global sentinels’; specifically, determining what regional AMR trends and emerging threats could be identified from whole genome sequencing of travel-acquired bacteria, and what this information can tell us about local microbial reservoirs. Our findings revealed geographic patterns of AMR gene prevalence, associations between traveler behavior and presence of virulence factors, and highlighted the role of plasmids in disseminating risk-associated genes both locally and globally. Our study demonstrates that traveler genomic surveillance could complement existing efforts to monitor the global prevalence and spread of AMR.

## Results

We recruited 608 travelers at US travel clinics between 2017 and 2020, collecting stool samples before and after international travel (*Methods*). Stool samples were screened for three target AMROs: ESBL-PE, CRE and mcr-E. As described in our previous study (29), 6.6% (40/608) of travelers were colonized with an AMRO before travel, and 38% (217/568) of those not colonized prior to travel returned with at least one AMRO, most often an ESBL-PE (98%, 212/217) (**Figure 1**). There was considerable regional variability in AMRO acquisition rates across the six travel regions investigated. Here, we sequenced 394 AMROs from 203 participants from this study, including from 186/217 (86%) of travelers with travel-acquired organisms. To capture multi-strain acquisition, we sequenced multiple strains from single stool samples where morphologically distinct organisms were identified in growth on selective media (66 stool samples had more than one distinct sequenced isolate; *Methods*). To avoid overrepresentation of acquired strains, we filtered out 86 isolates which were highly similar to others collected from the same individual (below a 50 SNP cutoff; Methods), leaving a total of 307 isolates for analysis. Long read sequence data were additionally generated for the first 216 (70%) isolates collected. The majority of isolates (292/307, 95%) were *E. coli*, and most were ESBL-PE (285/307; 92.8%) (**Table 1, Supplementary Data**).

**Figure 1.**
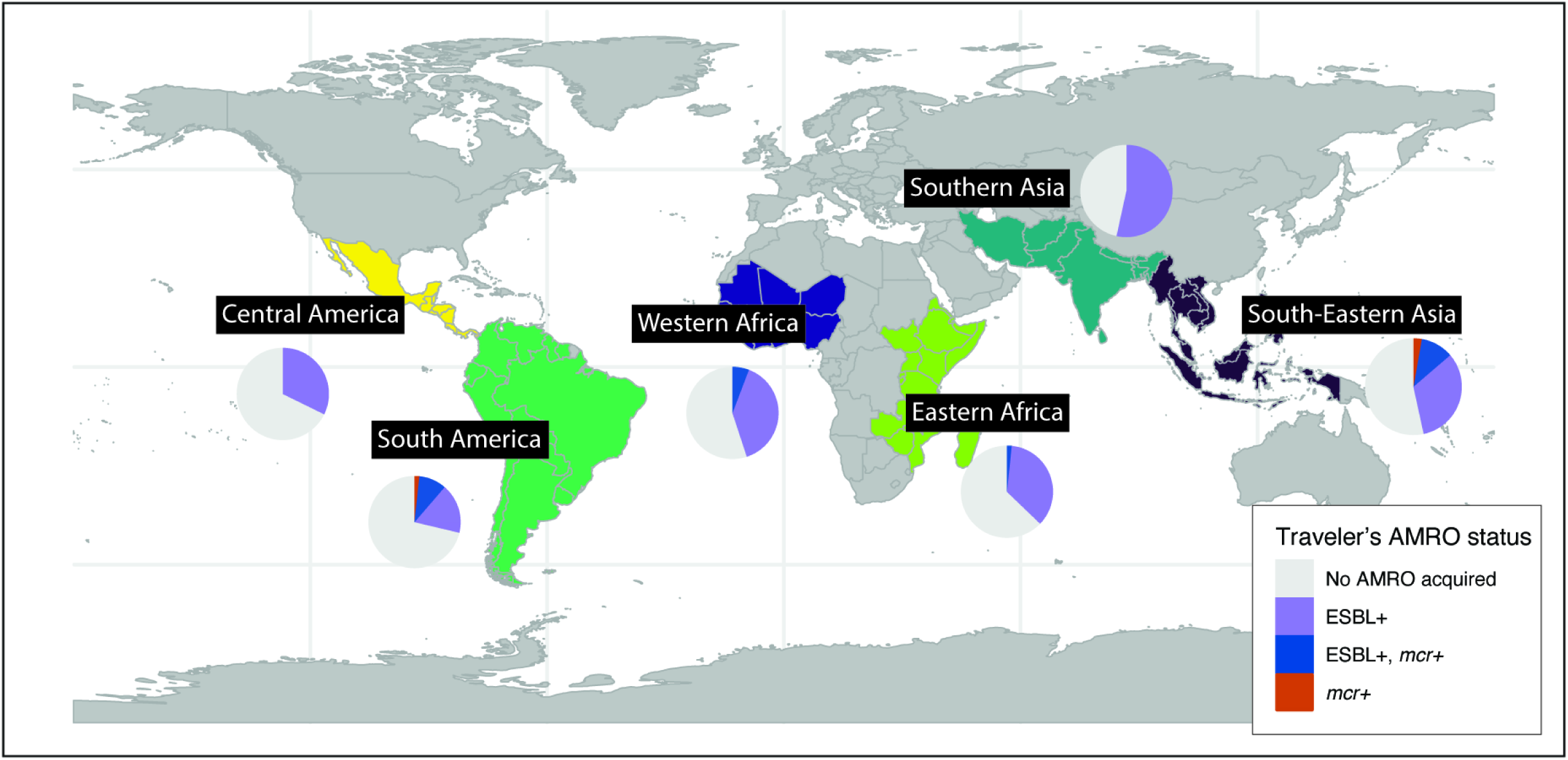
Global acquisition rates of AMR organisms. For the six most common travel regions, pie charts represent the proportions of travelers who returned carrying ESBL-PE and/or mcr-E. Regions are defined by the United Nations Statistics Division (*Methods*), and acquisition rates are calculated for the entire cohort of 608 travelers.

**Table 1.** Summary of traveler isolates.

|  | Count | % |
| --- | --- | --- |
| **AMRO (n=307)** |  |  |
| ESBL-PE | 285 | 92.8% |
| MCR-E | 28 | 9.1% |
| CRE | 2 | 0.7% |
| **Species (n=307)** |  |  |
| *E. coli* | 292 | 95.1% |
| *K. pneumoniae* | 10 | 3.3% |
| Other | 5 | 1.6% |
| **Region (n=307)** |  |  |
| Pre-travel | 45 | 14.7% |
| Southern Asia | 53 | 17.3% |
| South America | 45 | 14.7% |
| South-Eastern Asia | 43 | 14.0% |
| Eastern Africa | 42 | 13.7% |
| Western Africa | 38 | 12.4% |
| Central America | 13 | 4.2% |
| Other | 28 | 9.1% |
| **ESBL resistance mechanisms (n=285)** |  |  |
| CTX-M-15 | 177 | 62.1% |
| CTX-M-55 | 32 | 11.2% |
| CTX-M-27 | 44 | 15.4% |
| CTX-M-14 | 16 | 5.6% |
| Other CTX-M | 9 | 3.2% |
| Other ESBL | 10 | 3.5% |
| **Colistin resistance mechanisms (n=28)** |  |  |
| MCR-1 | 26 | 92.9% |
| MCR-3 | 3 | 10.7% |
| **CRE resistance mechanisms (n=2)** |  |  |
| NDM-5 | 2 | 100.0% |

### Travel-acquired AMR organisms are phylogenetically distinct from pre-travel strains

We first examined phylogenetic diversity of isolates belonging to the most common species, *E. coli*, to assess whether travel-acquired organisms reflected regional differences in strain type prevalence. Based on phylogroup, travel-acquired isolates were distinct from isolates carried pre-travel (p=5.3e-9, Chi-squared test); in particular, phylogroup B2 represented 43% (17/40) of pre-travel isolates, but only 7% (18/252) of travel-acquired isolates (**Figure 2**). Sequence type (ST) 131 within phylogroup B2 was overrepresented in pre-travel strains (OR=5.9, 95% CI: 2.3, 14.9; p=8e-5). Phylogroups A and B1 predominated post-travel (162/252; 64%). There were few differences in phylogroup distribution between travel regions among the post-travel isolates. While we observed that phylogroup D *E. coli* were slightly more common among isolates associated with travel to South-Eastern Asia (11/43, 26% vs. 11% in other travel regions; OR=3.1 (1.2, 7.4); Figure 2B), this was not significant after multiple testing correction (false discovery rate (FDR)=0.24)). Travel associated isolates exhibited considerable ST diversity; none of the three STs with more than 10 isolates were significantly overrepresented in any travel region.

**Figure 2.**
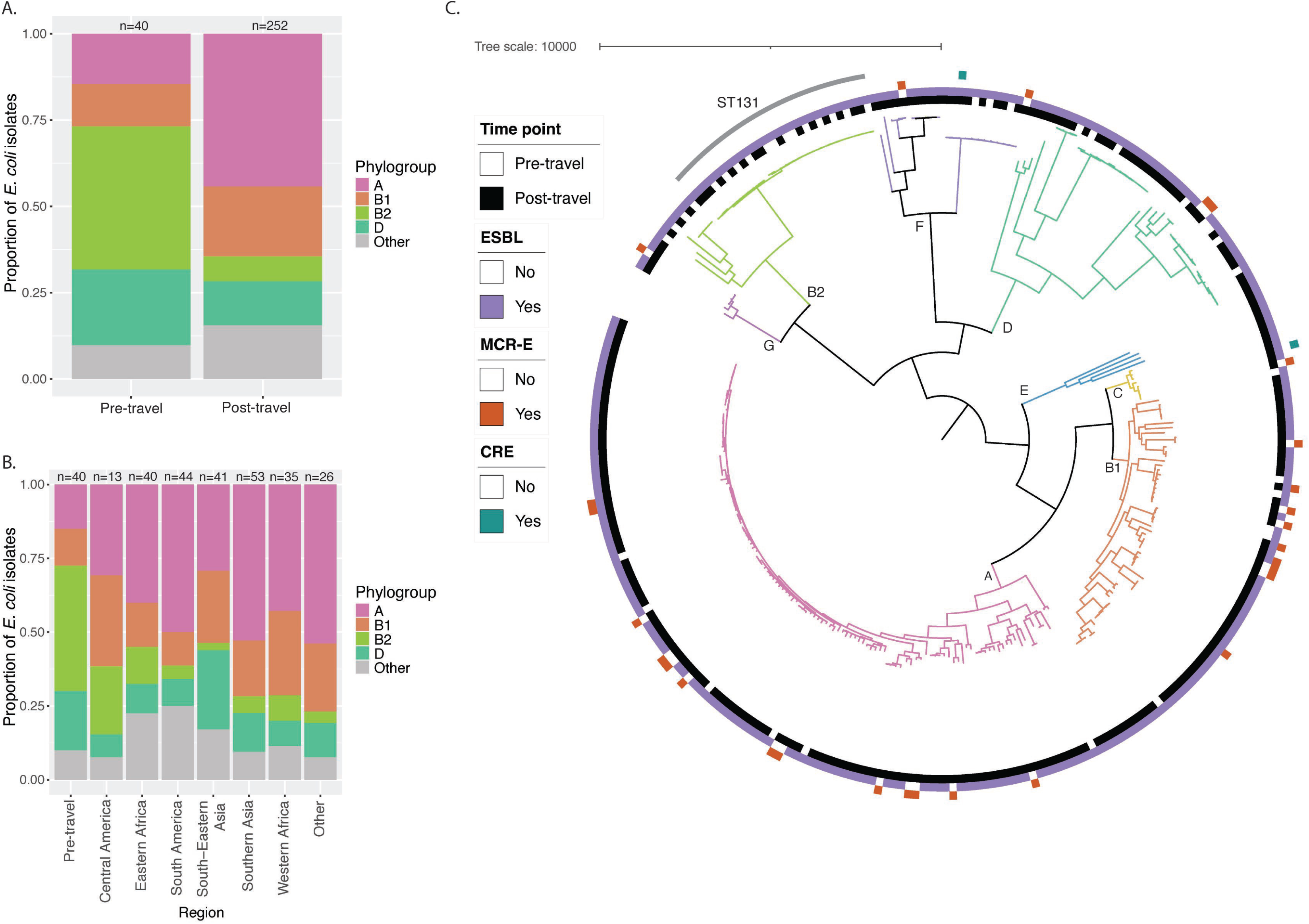
Phylogenetic distribution of acquired AMR *E. coli*. The phylogroup designation for all 292 *E. coli* isolates is compared (a) between pre-travel and post-travel, and (b) between each of the most common travel regions. (c) The phylogenetic tree is shown for all 292 *E. coli* isolates. Branches are colored and labeled by phylogroup.

### Acquired ESBL resistance mechanisms vary by travel region and environmental exposures

Although we observed little phylogenetic clustering by travel region, suggesting a limited role of specific bacterial lineages in driving local AMR prevalences, we hypothesized that horizontal gene transfer in *E. coli* might allow AMR genes to proliferate regionally on diverse genetic backgrounds. We therefore evaluated the resistance mechanisms associated with the observed resistance phenotypes to identify any geographic associations. Most ESBL-PEs carried a *bla*_CTX-M_ gene, including 96% (43/45) of pre-travel isolates and 97% (232/240) of post-travel isolates (**Figure 3a**). The most common allele in both pre- and post-travel samples was *bla*_CTX-M-15_, though the CTX-M allele distribution varied by geography. Compared to other travel regions, *bla*_CTX-M-27_ was significantly more common in isolates associated with South-Eastern Asia (OR=4.3 (95% CI: 1.7, 10.6); FDR=0.0034). *bla*_CTX-M-55_ was observed more frequently in isolates associated with South America (OR=7.9 (95% CI: 3.2, 19.7), FDR=2.8e-5) and South-Eastern Asia (OR=3.0 (95% CI: 1.1, 7.7), FDR=0.06), though the latter did not reach significance after multiple testing correction (**Figure 3b**). These findings are concordant with previous studies describing the expansion of *bla*_CTX-M-27_ in South-Eastern and Eastern Asia (29, 30) and the global prevalence of *bla*_CTX-M-55_ (31). There were ten non-*bla*_CTX-M_ ESBL-PE isolates, seven of which were collected from travelers returning from India. Resistance mechanisms in these isolates varied, including *bla*_CMY_ (n=5) and *bla*_SHV_ (n=3). Two unrelated *E. coli* isolates had no identified ESBL gene; resistance in these isolates was potentially mediated by *ampC* mutations. Two travelers acquired CRE during travel after visiting South-Eastern Asia (phylogroup C *E. coli*) and Southern Asia (phylogroup F *E. coli*); both isolates carried *bla*_NDM-5_ and *bla*_CTX-M-15_.

**Figure 3.**
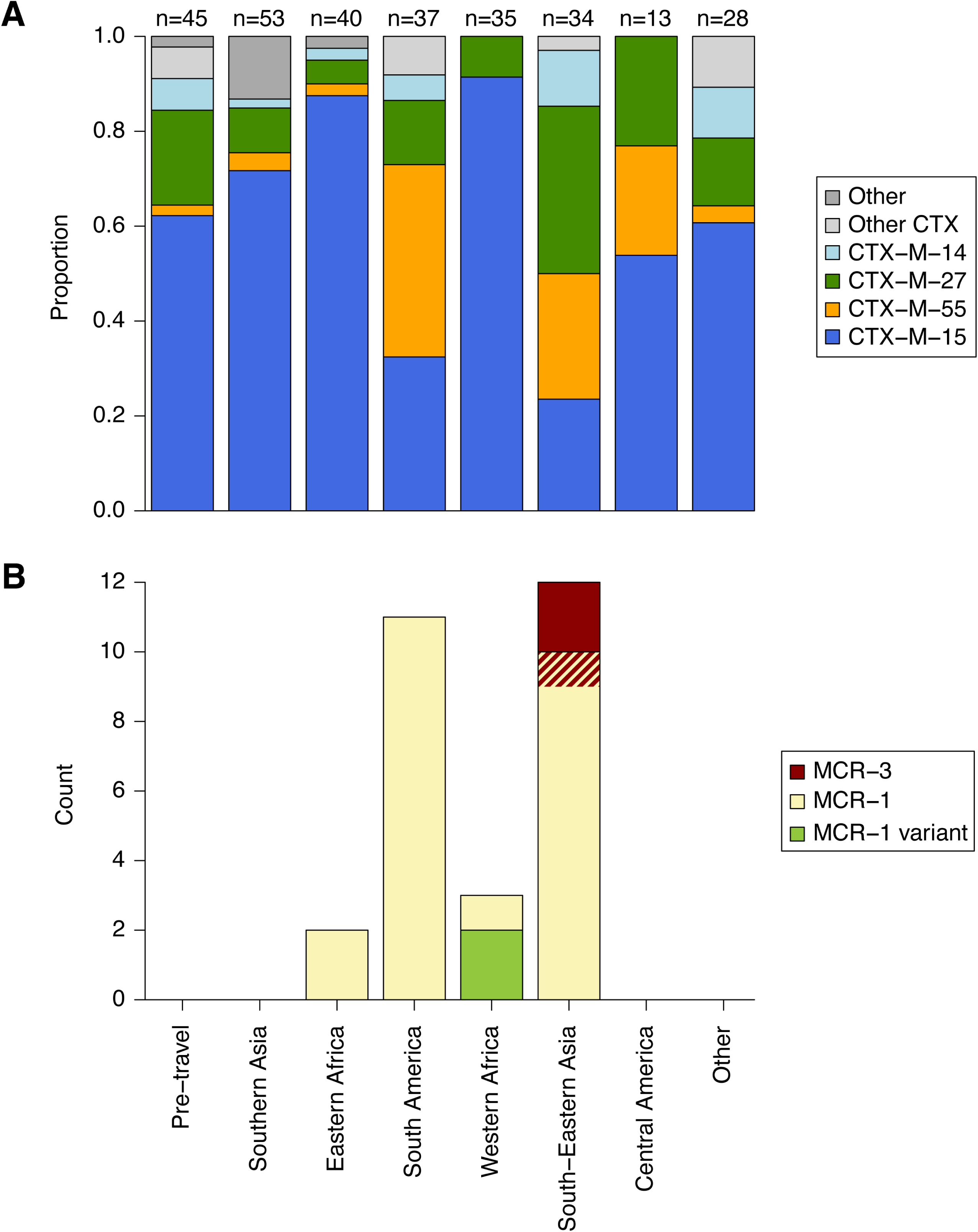
Resistance alleles show geographic heterogeneity. Genes mediating the observed resistance phenotypes are broken down by travel region. (a) Relative proportion of ESBL genes by region. The four most common CTX-M alleles are shown. Total counts are given above bars; counts exceed the number of ESBL organisms due to multiple ESBL gene carriage in 3 isolates (Supplementary Data). (b) *mcr* gene counts by region. Hatched region indicates one isolate carrying both *mcr-1* and *mcr-3*. *mcr-1* variants differ from *mcr-1.1* by a single mutation.

We next looked for positive associations between self-reported traveler behavior (see questionnaire, Supplemental Text) and ESBL gene carriage in post-travel isolates to assess potential sources of acquisition. Isolates collected from travelers who reported contact with animals, eating raw or undercooked fish, and eating food from a street vendor were significantly more likely to carry *bla*_CTX-M-55_, but none of these behaviors were significantly associated after adjusting for travel destination (**Table 2**). Isolates collected from travelers who reported drinking unpurified water were significantly more likely to carry *bla*_CTX-M-27_; this remained significant in multivariable regression after adjusting for travel destination (OR=3.33, 95% CI: 1.48, 7.44; p=0.0034).

**Table 2.** ESBL genes associated with traveler activities. ESBL genes with significant univariable associations to traveler activities are listed with estimated odds ratios (ORs), 95% confidence intervals (CIs) and p-values. Only travel-acquired ESBL isolates are considered (n=240). Regression results are also provided for models adjusting for travel destination.

| Gene | Traveler activity | Gene present (%) | Gene absent (%) | Univariable |  |  | Adjusted for travel destination |  |  |
| --- | --- | --- | --- | --- | --- | --- | --- | --- | --- |
|  |  |  |  | OR | 95% CI | p | OR | 95% CI | p |
| CTX-M-27 | Drinking unpurified tapwater | 15 (30%) | 35 (70%) | 3.64 | (1.68, 7.80) | 0.00088 | 3.33 | (1.48, 7.44) | 0.0034 |
| CTX-M-15 | Business travel |  |  | 2.42 | (1.22, 5.08) | 0.015 | 1.87 | (0.84, 4.34) | 0.13 |
| CTX-M-55 | Eating street food | 37 (50%) | 37 (50%) | 3.24 | (1.50, 7.10) | 0.0028 | 2.11 | (0.85, 5.25) | 0.12 |
| CTX-M-55 | Animal contact | 20 (20%) | 78 (80%) | 3.05 | (1.41, 6.92) | 0.0055 | 1.69 | (0.71, 4.14) | 0.24 |
| CTX-M-55 | Eating undercooked fish | 9 (43%) | 12 (57%) | 6.72 | (2.50, 17.74) | 0.00012 | 2.41 | (0.80, 7.07) | 0.14 |

In concordance with its known carriage on plasmids (31), *bla*_CTX-M-55_ was distributed broadly across the *E. coli* tree; at most, three *bla*_CTX-M-55_-carrying strains clustered within a common sequence type. In contrast, almost half (13/27; 48%) of *bla*_CTX-M-27_-carrying strains belonged to ST131 (OR=7.3 (95% CI: 2.9, 18.7); p=9.3e-6), corresponding to clonal spread of a known epidemic subclade (32). Of note, the plurality of *bla*_CTX-M-27_-carrying strains (6/13) were isolated from pre-travel samples.

### Diverse plasmids drive ESBL dissemination

Plasmids are important drivers of AMR dissemination, which is concordant with the minimal phylogenetic association between *E. coli* strains and ESBL alleles observed here. To explore the role of plasmids and better characterize the genomic neighborhood of ESBL genes, we generated near complete hybrid assemblies for the 216 isolates with long read sequence data available (*Methods*). All scaffolds were classified with MOB-Suite (33) to link genes to their host plasmid or chromosome. We found considerable diversity in the plasmid types associated with ESBL genes. For instance, *bla*_CTX-M-15_ was carried on plasmids spanning at least 18 distinct MOB-Suite clusters. There was limited association between the AMR gene host plasmid group and travel region (**Figure S1a**). Chromosomal carriage was more common for *bla*_CTX-M-14_ (43%) and *bla*_CTX-M-15_ (27%), and chromosomal integration of these genes was observed across all phylogroups (**Figure S1b**).

Given the association we observed between *bla*_CTX-M-55_ carriage and travel to both South America and South-Eastern Asia, we sought to determine whether horizontal transfer of the same genomic sequence could be driving its circulation in these specific global regions. Our analysis of the genomic neighborhoods surrounding each *bla*_CTX-M-55_ revealed a geographic pattern; the *bla*_CTX-M-55_ neighborhood in isolates brought back by travelers from South-Eastern Asia was commonly ISEcp1-hp-*bla*_CTX-M-55_-hp-Tn2 (corresponding to ‘Type I’ in a previous study(31), n=5/8) while isolates from travelers to South America typically followed the motif IS15-*bla*_CTX-M-55_-hp-(*bla*_TEM_)-IS26 (‘Type II’, n=6/7) (**Figure S2**). Previous analyses of *bla*_CTX-M-55_ contexts have shown type II to be increasing in prevalence, more common in genomes from South America, and often associated with animal sources (31).

### Colistin resistance gene variants and geographically localized plasmids identified among traveler samples

While the majority of the targeted AMROs identified from travelers were ESBL-PE, 28 isolates were mcr-E. All were *E. coli* from post-travel samples, most often associated with travel to South America (n=11) and South-Eastern Asia (n=12). Three isolates carried *mcr-3*, all associated with travel to South-Eastern Asia. Twenty-six carried *mcr-1*, with one carrying both *mcr-1* and *mcr-3* (**Figure 3b**). Most travelers colonized with mcr-E post-travel were also positive for ESBL-PE (18/23; 78%), either through acquisition of multi-resistant strains exhibiting both phenotypes (n=13), or co-carriage of multiple distinct *E. coli* strains (n=5). Two isolates associated with travel to Western Africa (Liberia and Ghana) carried an *mcr-1.1* gene with point mutations in the start codon, with retained functionality likely due to a subsequent second start codon. While start codon mutations have previously been observed in *mcr-1* (34), the specific mutations observed here appeared to be rare. The variant identified in a traveler returning from Liberia (M1R) had 13 perfect matches in the NCBI nt database (as of September 2023), two collected prior to 2020, while the variant identified in a traveler returning from Ghana (M1I) had no matches.

Two travelers visiting Peru in April and September 2018 returned with colistin-resistant *E. coli* strains, harboring highly similar IncI2 plasmids carrying *mcr-1*. The 60kb and 61kb plasmids shared 59kb at 99.8% nucleotide identity with two indels. Using BLAST to search for similar sequences to the longer plasmid in public databases, we found three other plasmids with query coverage and nucleotide identity at least as high as our identified pair. All three also originated from South America (Peru, Bolivia and Ecuador), had various bacterial hosts (*K. pneumoniae*, *Citrobacter braakii*, *E. coli*), and were isolated from human urine, food, and chicken, respectively.

### Regional variability in additional resistance genes

While we selected for ESBL-PE, CRE and mcr-E organisms in our study design, we sought to characterize additional phenotypic and genotypic features of all 307 AMRO isolates to assess other clinically relevant traits that might be associated with travel destination. We first considered resistance to other antimicrobials; all isolates were subjected to antimicrobial susceptibility testing against a panel of 19 antibiotics (*Methods*). In addition to beta-lactams, ESBL-PE isolates exhibited non-susceptibility to additional antibiotics : most commonly tetracycline (68% of ESBL-PE isolates), trimethoprim-sulfamethoxazole (65%) and aztreonam (64%). There was some regional variability in phenotypic AMR; gentamicin resistance was elevated in isolates associated with South-Eastern Asia (OR=3.2, 95% CI: 1.3, 7.6, FDR=0.044). Among isolates associated with travel to Southern Asia, 70% were non-susceptible to levofloxacin, higher than among isolates associated with other regions (OR=2.5, 95% CI: 1.3, 5.0; FDR=0.046) (**Figure S3**).

We additionally explored the full genotypic resistance profile, using RGI and CARD (35) to identify resistance genes in all isolates (*Methods*). Several genes were enriched in specific travel regions **(Figure S4**), often concordant with observed phenotypic patterns. Mutations in GyrA were more frequent in isolates collected from travelers to Southern Asia (OR=3.0, 95% CI: 1.5, 6.0; FDR=0.028), while *aac*(*3*)*-IIa* was enriched in ESBL isolates from South-Eastern Asia (OR=5.2, 95% CI: 2.0, 13.2; FDR=0.0092), contributing to the elevated resistance to fluoroquinolones and gentamicin in those regions, respectively. *bla*_TEM-1_ (OR=2.9, 95% CI: 1.4, 6.5; FDR=0.035) and *qnrS1* (OR=3.1, 95% CI: 1.4, 6.8; FDR=0.025) were both enriched in isolates associated with Western Africa. The genes *floR* (32.4% vs. 3.9%; OR=11.8, 95% CI: 4.3, 32.9; FDR=7.4e-6) and *fosA3* (40.5% vs. 1.8%; OR=37.1, 95% CI: 11.5, 143.1; FDR=2.0e-10), conferring resistance to phenicol and fosfomycin, respectively, were elevated in ESBL-PE isolates from travelers returning from South America.

### Virulence factor carriage highlights associations with ecology and traveler behavior

As AMROs inhabit a range of environmental niches with differential selective pressures, we hypothesized that they may contain additional gene content providing insight into their acquisition. To assess this, we looked for enriched virulence gene carriage associated with both travel regions and traveler activities.

Given the importance of *E. coli* as a cause of diarrhea, we first considered virulence genes used as markers for diarrheagenic *E. coli* pathotypes (*Methods*). On this basis, most strains (70%; 208/292) were non-diarrheagenic; however, 7.5% (22/292) were enteropathogenic *E. coli* (EPEC), 10.6% (31/292) were enteroaggregative *E. coli* (EAEC) and 10.9% (32/293) were diffusely adherent *E. coli* (DAEC) (**Figure S5**). Most EAEC isolates were observed post-travel; 42% (13/31) of them were associated with travel to Southern Asia. While our previous study noted that acquisition of AMROs was associated with self-reported diarrhea during travel (29), the rate of diarrhea was not elevated among travelers acquiring any specific pathotype. Next, we broadened our search for any additional virulence and stress factors (including metal resistance genes) with geographical enrichment. The salmochelin siderophore system encoded by the gene cluster *iroBCDEN* was enriched in isolates associated with travel to South-Eastern Asia (iroB: OR=5.5 (95% CI: 1.8, 15.4), FDR=0.0045) and South America (iroB: OR=6.2 (95% CI: 2.1, 17.0), FDR=0.0026), while metal resistance genes (the *sil*, *pco*, and *ter* gene clusters, encoding resistance to silver, copper and tellurite, respectively) were enriched in isolates linked to South-Eastern Asia (silC: OR=6.1 (95% CI: 2.3, 15.9), FDR=0.0014; pcoE: OR=4.6 (95% CI: 1.5, 13.4), FDR=0.036; terD: OR=5.4 (95% CI: 1.7, 16.0), FDR=0.019) (**Figure S6**).

We next considered genes associated with travel activity, and found several virulence genes to be significantly more common in isolates collected from travelers eating street food and reporting contact with animals. These included *iroBCDEN*, as well as the aerobactin operon *iucABCD-iutA* (**Table S1**). The genes *cvaC* and *iss* were also significantly more common in isolates from travelers reporting animal contact, and together with *iroBCDEN*, are among the markers for the ColV plasmid, a driver of avian pathogenic *E. coli* (35–39). We used a previously described marker-based approach to define ColV plasmids in our isolate collection (*Methods*) and found fifteen isolates (4.9%) meeting these criteria. ColV was more frequent in isolates linked to South America and in isolates associated with animal contact (OR=3.59, 95% CI: 1.14, 11.09; p=0.025, and OR=5.68, 95% CI: 1.66, 26.09; p=0.011 respectively, multivariable regression), highlighting the connection between travelers and high-risk environmental reservoirs.

Having identified geographical enrichment in ColV plasmid carriage, we looked across all plasmid groups reconstructed with hybrid assembly and classified with MOB-Suite to determine if any other potentially high-risk plasmids were more common in specific regions. While most plasmid groups identified with MOB-Suite were found across several travel destinations, some exhibited geographical association, including plasmids belonging to group 430, which were significantly more common in isolates linked to South-Eastern Asia. These plasmids carried a median of 14 ARGs, including either *mcr-1* or *mcr-3* (**Figure 4**). Plasmid colocation of multiple resistance genes represents elevated risk of rapid dissemination of multidrug resistance; detection of such plasmids highlights the importance of long read sequencing.

**Figure 4.**
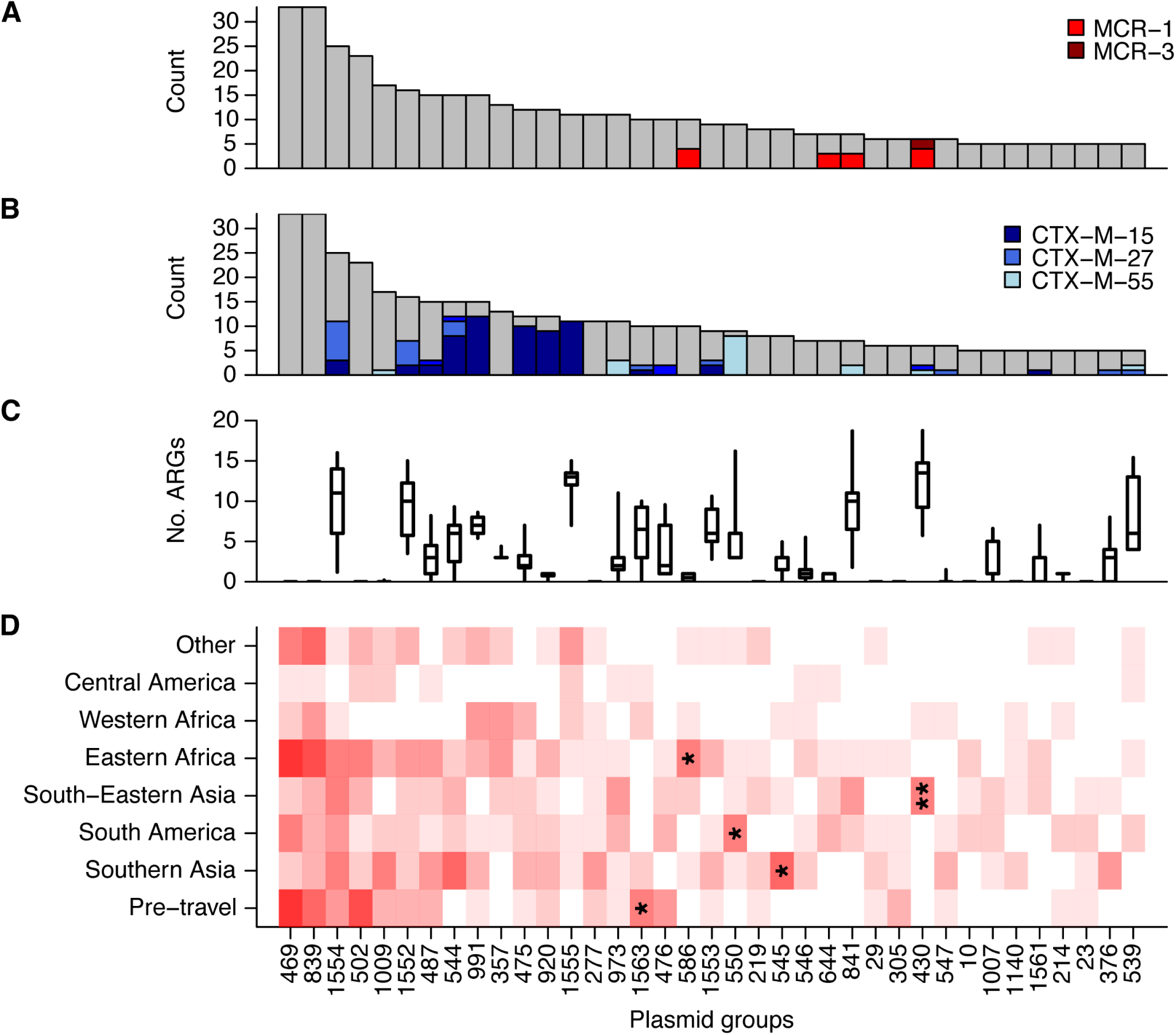
Plasmids vary in geographic distribution and ARG content. Plasmid content was identified with MOB-Suite for the 207 genomes with hybrid assemblies. (a) Frequency of the most common plasmid clusters across all isolates, with MCR gene content marked. (b) Frequency of the most common plasmid clusters across all isolates, with ESBL gene content marked. (c) ARG count distribution for each of the plasmid groups. Box plots display the interquartile range and median, as well as the 95% quantiles. (d) Heat map denoting frequency of plasmid group detection by travel region. Asterisks denote statistical significance after multiple testing correction; * FDR<0.05, ** FDR<0.005.

## Discussion

Robust surveillance and analysis of circulating AMROs across global regions is essential for mitigating the growing challenge of AMR. While considerable efforts are underway to establish global AMR surveillance networks (40), gaps remain, and implementing an international framework to collect appropriate samples is costly, resource intensive and presents numerous logistical challenges. In contrast, surveillance of international travelers requires only local sample collection and processing yet may provide a proxy insight into global AMR dynamics, complementing existing international surveillance efforts. In this study, we conducted genomic analyses on AMR bacterial isolates acquired by international travelers to explore global diversity and to identify regional patterns of resistance and virulence gene circulation, including potential emerging threats. We found that the majority of acquired AMROs were highly diverse *E. coli* with limited phylogeographic signal, though many AMR and virulence genes exhibited regional enrichment, driven by local plasmid-mediated dissemination. We additionally found associations between travel-related behavior and AMR gene carriage, suggestive of environment-specific bacterial and AMR reservoirs.

Several patterns identified in our study are concordant with previous analyses of global sample collections of *E. coli* from humans and wastewater, highlighting the utility of our indirect sampling of travelers. *E. coli* is known to be highly globally diverse, with limited geographic or environmental structure (41). While we found *E. coli* ST131 more frequently in pre-travel samples, travel-associated strains exhibited considerable diversity regardless of destination. *bla*_CTX-M_ alleles are known to vary by region, and their distribution has changed over time (30, 42), with new variants emerging (43). *bla*_CTX-M-55_ emerged in South-Eastern Asia (44), and was primarily found in Asia prior to 2017 (30, 44). We also found several examples in travelers returning from South America, indicative of the subsequent global dissemination of this gene observed as of 2022, particularly to the Americas (31). Characterizing the genomic neighborhood of this gene provides additional insight into regional circulation. While our sample size was limited, isolates associated with South America more frequently carried *bla*_CTX-M-55_ in a ‘type II’ background, which was more common in this region, and also linked to animal populations and food sources, based on a previous study of over 2,000 *bla*_CTX-M-55_ isolates (31).

Our sequencing-based approach further enabled us to find that silver and copper resistance genes were more common in isolates associated with travel to South-Eastern Asia. This could indicate increased exposure to industrial pollution (45, 46), although further work is needed to explore the correlations of metal resistance genes and environmental contamination. Finally, our finding that animal contact was associated with elevated levels of ColV plasmid carriage in acquired AMROs is striking, given its known high prevalence among animal sources (47), and highlights the potential for future traveler surveillance efforts to link acquired genes and their associated primary reservoirs.

Earlier studies of international travelers have used PCR-based methodology to examine global AMR patterns, finding similar trends in e.g. *bla*_CTX-M-15_ as the dominant ESBL gene acquired by travelers, with particularly high levels associated with travel to sub-Saharan Africa (14, 48). However, such approaches cannot provide further insight into gene context, co-carriage of additional virulence or resistance genes, or plasmid content. To our knowledge, this is the largest study to date utilizing whole genome sequencing of travel-acquired AMROs to describe AMR patterns associated with travel regions; previous sequencing studies did not attempt to identify geographic patterns due to sample size or sampling approach (18–20, 22–26). Hybrid sequencing further allowed us to associate key genes with their diverse plasmid backgrounds. The ability to characterize the genomic context around target genes allows one to track mobile sequences across strains and plasmids and identify associations with particular environmental niches (49). Through hybrid assembly, we identified frequent resistance gene co-location on plasmids, including an *mcr*-carrying plasmid group associated with South-Eastern Asia harboring an average of 14 additional ARGs. Such plasmids pose a significant public health concern, and their dissemination should be monitored.

Travelers acquire AMR organisms at vastly different rates depending on destination; however, the utility of travelers as sentinels to report on regional genomic landscapes has yet to be explored. Such a traveler-based approach could identify potential threats emerging in regions with limited surveillance. This is highlighted in our study with the detection of rare mutations in the *mcr-1* gene in two travelers returning from Western Africa. Tracking phylogeographic patterns of clinically important genes, including *mcr-1*, is essential both for monitoring dissemination pathways, and the ability to rapidly respond to emerging threats.

Our study has limitations. While our isolate collection was larger than any previous study of traveler AMROs, the sample size per travel region was relatively limited, hampering the ability to estimate prevalence of genomic features with precision. This is especially true for insights into plasmids, as long-read data was not generated for all isolates. Moreover, the aggregation of travel destinations into global regions may obscure important heterogeneity between and even within specific countries. Nevertheless, we were able to detect significant differences in resistance alleles and plasmids between regions, which could be explored at a more granular level in future studies or in ongoing surveillance. By picking morphologically distinct colonies from selective media, we captured greater diversity than a standard approach of taking a single isolate per individual, though we may still be missing additional, distinct strains. Further, although we filtered out highly similar same-host isolates to prevent overrepresentation of sequence types and genes, within-host horizontal gene transfer could nevertheless result in certain genes appearing more prolific. Since all studied isolates belonged to three specific AMR phenotypes, conclusions related to phylogenetic distributions, additional gene content and plasmids are not necessarily generalizable, due to colocation or interaction with targeted resistance genes.

While there have been efforts to expand global surveillance of AMR, including through multinational networks (50) and targeted community-level efforts (51–55), coverage remains sparse and requires extensive coordination to collect reliable samples. Given the nature of international travel, using travelers as sentinels could be an important approach to provide complementary AMR surveillance, including offering greater visibility in regions where other AMR surveillance approaches are limited. Such a framework would be ideally implemented prospectively to understand AMR trends in real time. Traveler pathogen genomic surveillance, if implemented systematically with geographic representation, could provide insight on global AMR dynamics, and importantly, provide information about global dissemination of emerging AMR threats.

## Methods

### Study design & sample collection

We recruited participants at five U.S. travel clinics affiliated with Global TravEpiNet (56), located in Boston MA, New York NY, Bronx NY, Salt Lake City UT and LeHigh Valley, PA, during a pre-travel health visit, as previously described (29). Participants were aged 1-85 years, and no exclusion criteria were applied. Recruitment began in 2017 at the Boston site, and in 2018 for all other sites, and concluded in 2020. Written informed consent was obtained from all participants in the study. Institutional review board approval was obtained from the human research committee at each participating enrollment site. Participants self-collected pre- and post-travel stool samples, which were immediately stored in Cary Blair medium and mailed to the Massachusetts General Hospital clinical microbiology laboratory. Benchmarking was performed to assess the recovery of AMR organisms under this sample collection protocol (57). Samples were screened for the presence of extended spectrum beta-lactamase producing Enterobacterales (ESBL-PE), *mcr*-mediated colistin-resistant Enterobacterales (mcr-E) and carbapenem-resistant Enterobacterales (CRE) as previously described (57, 58). Briefly, cultures were screened for ESBL-PE by growth on HardyCHROM ESBL (Hardy Diagnostics, Santa Maria, CA) medium at 35°C and for CRE by either growth on McConkey agar following incubation in tryptic soy broth inoculated with a 10-µg meropenem disc ((59)) or growth on CHROMID CARBA (bioMerieux, Durham, NC) medium at 35°C. Colistin resistance was screened for using Luria Bertani agar containing colistin sulfate, vancomycin, and amphotericin (58). Resistant colonies were picked from the selective screening; multiple colonies were picked if there were distinct morphotypes. Recovered organisms underwent initial antimicrobial susceptibility testing (AST) against a panel of 18 antibiotics using the Vitek2 antimicrobial susceptibility testing automated system (bioMérieux, Durham, NC); ampicillin, amoxicillin, ampicillin/sulbactam, piperacillin, cefazolin, ceftriaxone, cefepime, aztreonam, ertapenem, imipenem, meropenem, amikacin, gentamicin, ciprofloxacin, levofloxacin, tetracycline, nitrofurantoin, trimethoprim-sulfamethoxazole. Colistin AST was also performed using an internally validated broth microdilution panel (Sensititre, ThermoFisher Scientific, Waltham, MA). Ceftriaxone, carbapenem, and colistin non-susceptibility was confirmed using phenotypic (ESBL-PE, CRE) or molecular (mcr-E) testing as previously described (58). Healthcare providers used structured questionnaires before travel to collect information on demographics, health, travel itineraries and activities, medications and symptoms. Travelers completed a questionnaire after travel regarding behaviors and illness while traveling (see traveler questionnaire; Supplemental Text).

### Whole genome sequencing

#### DNA extraction

1 ml of a 4 ml overnight culture of each isolate grown in Luria Bertani broth was centrifuged for 10 minutes at 7500 rpm to pellet cells. The supernatant was discarded and pellets were processed using the Qiagen QIAamp DNA Mini kit. The only deviations from the standard protocol were that samples were not incubated at 70°C following addition of Buffer AL, and samples were eluted in 100 µl distilled water. DNA concentrations (ng/µl) were measured using a Nanodrop.

#### Illumina sequencing

Illumina whole genome paired-end libraries were prepared for a total of 393 isolates as previously described, and sequenced on Illumina HiSeq 2500 or HiSeq X sequencers at the Broad Institute (60).

#### Oxford Nanopore sequencing

600 ng of DNA from each sample was used as input into the Oxford Nanopore 1D ligation library construction protocol (SQK-LSK109) following the manufacturer’s recommendation. Samples were barcoded using the Native Barcoding Expansion 1-12 kit to run in batches of between 1 and 10 samples per flow cell on a GridIon (Oxford Nanopore Technologies Ltd, Science Park, UK). Samples were run on flowcells FLO-MIN106D (R9.4.1) and base-called using ont-guppy-for-minknow v2.0.5. Due to resource limitations, ONT data were generated for 263 isolates, approximately representing the first two-thirds of isolates received.

### Sequence data analysis

#### Assembly and annotation

All Illumina short read datasets were assembled using Spades v3.11.1 (61), and annotated with Prokka v1.14.6 (62). For the subset of isolates with long read data available, genomes were assembled using UniCycler v0.4.4 (63) pipeline described previously (49), and annotated using the Broad Institute’s prokaryotic annotation pipeline, as previously described (64).

#### Removal of duplicate isolates

While colonies were selected based visually on differential morphology, some sequenced isolates from the same sample were nearly identical. To avoid overrepresentation of strains with a recent common ancestor which may represent the same exposure event, we used read-based alignment against the GenBank reference genome CP015159.1 to calculate pairwise SNP distances between all isolates. Strain pairs from the same individual separated by <50 SNPs were considered to belong to the same lineage, and one isolate was removed from further consideration. If ONT data were available for one isolate, this was retained; if isolates represented both pre- and post-travel, the pre-travel isolate was retained; otherwise, one isolate was retained at random. A total of 86 isolates were removed on the basis of this filtering, leaving 307 isolates for analysis. Post filtering, seven pairs of isolates collected from the same individual belonged to the same sequence type; the pairwise distances ranged from 500-12,000 SNPs, and most exhibited distinct antimicrobial susceptibility profiles. These isolates were therefore considered distinct and were retained for analysis.

#### Gene identification and pathotyping

Resistance genes were identified using CARD and RGI v5.5.1 (35). Virulence and stress factors were identified using AMRFinderPlus v3.11.2 (65). Pathotypes were determined based on presence of pathotype-associated virulence genes; these were determined to be those most commonly used for pathotype classification in the literature (66–68). Enteropathogenic *E. coli* (EPEC) were defined as isolates containing genes *eae* or *bfp*; enteroaggregative *E. coli* (EAEC) were those containing *agg*, *aat*, or *aai*; diffusely adherent *E. coli* (DAEC) were those containing *afa* or *dra*; enteroinvasive *E. coli* (EIEC) were those containing *ipa*; enterotoxigenic *E. coli* (ETEC) were those containing *elt* or *est*; and enterohaemorrhagic *E. coli* (EHEC) were those containing *stx*. Any isolates not containing any of these genes were defined as non-diarrheagenic (non-DEC) *E. coli*.

As previously defined (39), we considered an isolate ColV-positive if it carried at least one or more genes from four or more of the following six gene sets (i) *cvaABC* and *cvi* (the ColV operon), (ii) *iroBCDEN* (salmochelin), (iii) *iucABCD* and *iutA* (aerobactin), (iv) *etsABC*, (v) *ompT* and *hlyF*, and (vi) *sitABCD*. Since not all of these genes belong to the AMRFinderPlus database, we used BLAST to identify relevant hits to the Virulence Factor Database (69).

#### Phylogenetic tree construction

Core genome alignment was performed using Parsnp (Treangen et al. 2014) on all 292 *E. coli* isolate assemblies. Isolate GTEN_1 was used as the arbitrary ‘reference’ genome. Recombination removal and tree construction was performed using Gubbins (70) with RAxML (71). The *E. coli* phylogeny was visualized in iTOL (72) using midpoint rooting.

#### Plasmid classification

Plasmid content within the hybrid assemblies was classified using MOB-Suite version v1.4.9 (33). Plasmids were assessed for pairwise shared, contiguous content using ConSequences (49).

### Statistical analysis

Travel destination was provided at country level; to increase per-destination sample size, we aggregated travel destinations into geographic regions according to the United Nations Statistics Division M49 standard (subregion level) (73). We grouped regions with fewer than 10 travelers together into an ‘Other’ category. To test for phylogroup heterogeneity between pre- and post-travel isolates, we considered the four most common phylogroups (A, B1, B2, D) and applied the chi-squared test. Associations between binary features (e.g. gene presence, plasmid presence, antibiotic non-susceptibility) and geographic region were assessed using Fisher’s exact test; odds ratios, confidence intervals and p-values were computed in R v4.2.1. Multiple testing correction was performed using the Benjamini Hochberg procedure. Multivariable logistic regression models were used to assess the roles of travel destination and travel activity on gene and plasmid presence, and were fit using the glm function in R in the form `glm(gene presence∼activity+destination, family=”binomial”)`.

## Data availability

Sequence data for this project is available in SRA under Bioproject PRJNA528511. Accession numbers are provided in Supplementary Data isolate_summary_table.

## Acknowledgements

We thank members of the Broad Bacterial Genomics group for helpful discussions. We also thank members of the Broad Institute Microbial ‘Omics Core’ and Genomics Platform for their assistance with data generation.

